# Phenotypic plasticity, stalk geometry, and noncoding variation underpin stalk lodging resistance in maize

**DOI:** 10.64898/2026.08.24.746877

**Authors:** Bharath Kunduru, Norbert T. Bokros, Kaitlin Tabaracci, Rohit Kumar, Manwinder S. Brar, Christopher J. Stubbs, Yusuf Oduntan, Caique Machado e Silva, William C. Bridges, Ravi V. Mural, Seth DeBolt, Gota Morota, Christopher S. McMahan, Daniel J. Robertson, Rajandeep S. Sekhon

## Abstract

Stalk lodging causes severe yield losses in maize (*Zea mays* L.) worldwide, worsening food and feed security. Stalk lodging resistance is influenced by multiple traits at various levels of biological organization, collectively referred to as intermediate traits, but their identities, genetic bases, and interrelationships remain poorly resolved. Here, evaluation of multiple geometric and structural intermediate traits in a maize diversity panel across four environments showed that macroenvironmental variation is the predominant driver of phenotype plasticity and that plasticity varies with internode position along the stalk, consistent with height-dependent mechanosensing. Major and minor diameters, moment of inertia, and rind penetration resistance, were genetically tractable and showed strong genetic correlations with stalk flexural stiffness. Multivariate analyses revealed two distinct but complementary mechanistic pathways, represented by cross-sectional geometry and rind architecture, that contribute to stalk mechanical performance. Association analyses using whole-genome resequencing data identified 705 SNPs associated with intermediate traits, fewer than 20% of which overlapped genic regions, indicating that most associated variation resides outside annotated genes. Interestingly, about 22% of SNPs were shared between at least two traits, indicating substantial shared genetic control among intermediate traits. Candidate gene analyses highlighted novel promising candidate loci associated with intermediate traits while recovering genes previously implicated in stalk lodging resistance. The predominance of noncoding associations further suggests that regulatory variation may contribute substantially to natural variation in intermediate traits underlying stalk lodging resistance.

**Box 1 – Description of key terms:** *Stalk lodging resistance:* A conceptual, holistic assessment of the ability of a genotype to withstand external forces, including wind and gravity, and biotic factors, including insect pests and diseases that contribute to stalk lodging.

*Lodging incidence:* A count or proportion of plants lodged in a defined area.

*Stalk flexural stiffness (a.k.a. stalk flexural rigidity):* A measure of the ability of a stalk to resist bending deformation such that higher flexural stiffness indicates a higher magnitude of the force required to bend a stalk to a certain distance.

*Stalk bending strength:* The maximum bending moment a stalk can support before undergoing permanent mechanical deformation.

*Structural property:* Characteristics of a structure that affect its mechanical response to physical loads and are the resultant combination of geometric and material properties.

*Geometric property:* The property of an object derived from its geometric form, including size, shape, length, thickness, curvature, etc.

*Macroenvironment:* Broad set of external factors, including meteorological conditions, soil properties, topography, biological environment, etc., that collectively influence the growth, development, and performance of a genotype across different geographical locations.

*Microenvironment:* Local physical, chemical, and biological factors in the immediate vicinity of a genotype that influence growth, development, and performance.

## Introduction

Stalk lodging, the permanent angular displacement of plant stems, remains a persistent and serious challenge in cereal crop production. Rising global demand for cereals has driven both genetic improvement and the adoption of modern agronomic practices, including high-input fertilization and increased planting densities aimed at maximizing yield per unit area. While these strategies have effectively boosted productivity, an unintended consequence has been an increased incidence of stalk lodging. In maize (*Zea mays* L.), one of the most widely grown cereal crops globally and a vital source of food, feed, and industrial products, lodging can cause over 20% losses in grain yield. Based on the survey of yield losses reported during weather events and recent trends in global maize production and trade (Elmore and Ferguson, 1999; FAO, 2024; USDA, 2026), preventing just 1% of stalk lodging could save the global maize industry more than $3 billion annually.

Stalk lodging reduces yield and crop quality through multiple, interconnected physiological and agronomic mechanisms. It disrupts photosynthesis and impairs vascular transport of photosynthates, water, and nutrients, thereby limiting grain filling and reducing overall productivity. Lodged stalks also hinder mechanical harvesting and lower grain quality by exposing ears to soil moisture, which can lead to kernel imbibition and premature germination. Furthermore, breakage sites provide entry points for pests and stalk rot pathogens, accelerating tissue degradation and diminishing stover quality. These combined effects underscore the importance of identifying the phenotypic and genetic determinants of stalk integrity under modern agricultural conditions.

Stalk lodging resistance, the ability of plant stalks to withstand lodging-inducing forces, is a complex trait determined by the interaction between stalk physical properties and environmental conditions (Sekhon et al., 2020; Xue et al., 2020; Bokros et al., 2024). However, the specific stalk traits and environmental drivers underlying lodging resistance are poorly understood, impeding progress in breeding programs focused on higher yield with superior lodging resistance (Robertson et al., 2017; Huang et al., 2025). Stalk lodging resistance is determined by a combination of structural, geometric, and material properties, collectively referred to as intermediate traits, that shape stalk structural performance, although the identity and functional relevance of many such traits remain unresolved (Kunduru et al., 2023; Stubbs et al., 2023). While certain intermediate traits such as plant height, rind thickness, and cell wall composition have been studied and selectively incorporated into breeding programs (Sekhon et al., 2020; Stubbs et al., 2020), most intermediate traits still lack well-defined genetic underpinnings. Among these, reduced plant height has been widely exploited to improve lodging resistance across major cereal crops, including maize (Kuczyńska et al., 2013; Zhao et al., 2022). Short-stature maize hybrids are less susceptible to stalk lodging, exhibit greater dry matter partitioning to grain, and provide opportunities to optimize crop management practices while maintaining yields comparable to those of traditional tall hybrids (Barten et al., 2022; Kosola et al., 2023). However, the increased susceptibility of plants to stalk lodging under high nitrogen fertilizer inputs and dense planting continues to constrain crop productivity in maize (Shah et al., 2021). Furthermore, dwarf genotypes are often susceptible to drought, highlighting the need to identify stalk attributes beyond short-stature to develop lodging-resistant genotypes with superior yield performance (Vikram et al., 2015; Jatayev et al., 2020).

Several studies have sought to elucidate the identity and genetic basis of intermediate traits associated with stalk lodging resistance in maize (Kumar et al., 2021; Wang et al., 2024). However, differences in phenotyping platforms and methodologies have limited comparability across studies and constrained translation of these findings into breeding programs. Recent advances in phenotyping have begun to address this gap. The development of the Device for Assessing Resistance to Lodging IN Grains (DARLING) enables field-based measurement of stalk bending strength and flexural stiffness under biologically relevant loading conditions, allowing high-throughput testing that closely mimics natural failure patterns (Cook et al., 2019). Stalk bending strength, a destructive parameter, and stalk flexural stiffness, a non-destructive parameter, were reported to be excellent indicators of natural stalk lodging incidence (Robertson et al., 2016; Sekhon et al., 2020). We developed a laboratory-based rind puncture technique that provides accurate, high-throughput measurements of rind thickness and diameter (Seegmiller et al., 2020). These tools now offer excellent opportunities for generating reproducible phenotypic data to support genetic studies of lodging resistance.

In this study, we evaluated stalk strength and a comprehensive suite of intermediate traits in a genetically diverse maize inbred panel grown across multiple environments. We leveraged field-deployable phenotyping devices designed to capture biologically meaningful mechanical traits and integrated these data with genome-wide marker information. Our objectives were to (1) characterize natural variation in lodging-related traits, (2) identify key traits predictive of stalk lodging resistance, and (3) dissect the genetic architecture underlying these traits. This work provides new insights into the biomechanical and genetic basis of stalk strength and lays the foundation for predictive genome-to-phenome modeling of complex agronomic traits.

## Materials and methods

### Plant material and experimental design

Details of plant material, experimental design, agronomic practices, weather data, and phenotype data corresponding to the present study are available elsewhere (Kunduru et al., 2025; Silva et al., 2025). Briefly, a maize diversity panel comprising 552 inbred lines was evaluated across four environments, defined by the combination of two geographic locations and two years (Table S1). The inbred panel represents eight genetically distinct classes of maize germplasm, including stiff stalk, non-stiff stalk, iodent, sweet corn, popcorn, and tropical genotypes. In each environment, the panel was planted in a randomized complete block design with two replications. Each plot consisted of a single 7.62 m row with 0.76 m row spacing, resulting in a total plot area of 5.8 m². Notably, not all 552 inbred lines were evaluated in every environment and, due to irregular germination and strict quality control during phenotypic data collection and post-processing, the number of inbred lines evaluated ranged from 363 to 495.

### Description of intermediate traits

We studied nine intermediate traits representing the geometric and structural properties associated with stalk lodging resistance. Geometric properties included plant height, ear height, major diameter, minor diameter, rind thickness, and moment of inertia, while the structural properties included stalk flexural stiffness, stalk bending strength, and rind penetration resistance. Stalk traits, including plant height, ear height, stalk flexural stiffness, and stalk bending strength, were recorded for each stalk, whereas internode traits, including major diameter, minor diameter, rind thickness, rind penetration resistance, and moment of inertia were measured for individual internodes. Internode traits were collected on five distinct internodes identified as the internode immediately below the primary ear-bearing node, hereafter the ear internode (E), and the first four elongated internodes starting at the bottommost elongated internode, hereafter the bottom internode (B1), labeled B1 through B4 (Figure S1). Phenotyping protocols of these intermediate traits are described in detail elsewhere (Kunduru et al., 2023; Tabaracci et al., 2024). Briefly, within each plot, ten healthy and representative plants were sampled and tagged with unique identifiers to preserve their identity and field coordinates throughout the phenotyping pipeline. This approach allowed for quality control during post-processing of the phenotypic data.

### Phenotypic data analysis

For each trait, the following linear mixed-effects model was fitted using the *lme4* (version 1.1.38) R package (Bates et al., 2015):

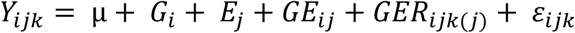

where *Yijk* denotes the raw phenotypic value for the inbred line *i* in the replication *k* nested within the environment *j*, µ is the grand mean across genotypes and environments, *Gi* is the fixed effect of the inbred line *i*, *Ej* is the random effect of the environment *j* and is ∼NID (0,σ*_E_*^2^), *GEij* is the random effect of the interaction of inbred line *i* with environment *j* and is ∼NID (0,σ*_GE_*^2^), *GERijk(j)* is the random effect of interaction of inbred line *i* with replication *k* nested within the environment *j* and is ∼NID (0,σ*_GER_*^2^), and *εijk* is the random residual error and is ∼NID (0,σ*_ε_*^2^). Estimated marginal means corresponding to the genotype effect (adjusted genotype means) for each inbred line were then obtained across environments using the *emmeans* (version 2.0.0) R package (Lenth and Piaskowski, 2025). These adjusted means were used to perform GWA analyses and also to estimate SNP heritability and genetic correlation through the genome-wide complex trait analysis tool (version 1.95.3) (Yang et al., 2011).

To estimate variance components for each trait, the same linear mixed-effects model was refitted using the *lme4* R package, with all model terms specified as random effects while retaining the intercept as the only fixed effect. When singular fits occurred, random-effect terms contributing negligible variance were sequentially removed until the model converged without singularity.

We fitted the following model to examine the phenotypic plasticity, defined as the ability of a genotype to produce distinct phenotypes in different environments, of the inbred lines (Bradshaw, 1965):

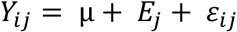

where *Yij* denotes the observed phenotypic value of the inbred line *i* in the environment *j*, *µ* is the grand mean across environments, *Ej* is the fixed effect of environment *j*, and *εij* is the random residual error and is ∼NID (0,σ*_ε_*^2^). Only those inbred lines evaluated in all four environments were included in the analysis. For each inbred line, we used the phenotypic values for individual traits measured in one environment as a baseline and compared the phenotypic values recorded in other three environments against this baseline to assess plasticity. P-values less than α = 0.05 were considered indicative of significant phenotypic plasticity. Within each environment, an unpaired two-sample Wilcoxon test was used to assess equality of replicate means, serving as an indicator of microenvironmental effects on intermediate traits.

Principal component analysis and phenotype correlations were performed using the *factoextra* (version 2.2.0) and *corrplot* (version 0.95) R packages, respectively (Wei and Simko, 2024; Kassambara and Mundt, 2026). We employed extreme gradient boosting (XGboost), a machine learning algorithm capable of capturing non-linear relationships and higher-order interactions, to model stalk structural traits (Chen and Guestrin, 2016). The methodology employed for this predictive modeling was discussed elsewhere (Kunduru et al., 2023). Pairwise comparisons of allelic effects for significant SNPs were performed using Wilcoxon rank-sum tests with Benjamini–Hochberg correction for multiple testing (Wickham et al., 2019). Computation and mapping of linkage disequilibrium between SNPs was performed using the *LDheatmap* (version 0.95) R package (Shin et al., 2006). R (version 4.3.1) was used to perform all analyses discussed in the present study (R Core Team, 2023).

### Genotype data and quality control

Whole genome resequencing data for the 552 inbred lines were downloaded from the Dryad database and processed following published pipelines (Grzybowski et al., 2023). Briefly, the sequence reads with an average sequence depth of 22x were aligned to the maize Zm-B73-REFERENCE-NAM 5.0 reference genome (Hufford et al., 2021). Variant filtering and quality control were performed with bcftools (version 1.19) (Li et al., 2009) and PLINK2 (version Alpha 4.3) (Chang et al., 2015). We excluded i) indels and multiallelic single nucleotide polymorphisms (SNPs), ii) individuals with more than 5% missing data, and iii) SNPs with minor allele frequency less than 0.05 and more than 5% missing data. The final genotype dataset contained 14,359,923 SNPs with an average genome-wide marker density of 6,736 SNPs per megabase (Mb). At the chromosome level, marker density ranged from 6,258 to 7,127 SNPs per Mb (Figure S2). The complete genotype matrix for 552 inbred lines was retained for all genome-wide association (GWA) analyses. However, each analysis included only the subset of lines with available adjusted phenotypic data for the corresponding trait.

### Genome-wide association analyses

To identify the marker-trait associations for intermediate traits, we performed bootstrap resampling association analyses with the FarmCPU model (parameter settings of maxLoop = 10 and method.bin = FaST-LMM) implemented using the *rMVP* (version 1.4.0) R package (Valdar et al., 2009; Yin et al., 2021). For each trait we executed 100 iterations, with each iteration analyzing 90% of the randomly sampled phenotype data. Within each iteration, SNPs were considered significant at a Bonferroni-corrected threshold (α = 0.05) based on the effective number of markers estimated using the genetic type I error calculator implemented in the KGGSEE tool (version 1.0) (Li et al., 2012). Each SNP was assigned a resample model inclusion probability (RMIP) score equivalent to the fraction of iterations in which it was significant. For downstream analyses, significant SNPs exhibiting RMIP scores of at least 0.05 were selected (Mural et al., 2022). To account for population structure, the kinship matrix was incorporated as a random effect, and the first five principal components were incorporated as covariates. The kinship matrix and principal components were calculated using the *MVP.K.VanRaden* and *MVP.PCA* functions available in the *rMVP* tool, respectively. All genes within ±10 kilobase (kb) window from the retained SNPs were considered as candidates for genetic analyses.

## Results

### Phenotype variation and plasticity for intermediate traits

The high-density phenotype dataset, comprising measurements on five internodes of 30,584 stalks across 552 maize inbred lines representing eight maize germplasm classes, revealed substantial variation for all intermediate traits studied (Figure 1A). Notably, all internode traits, including major diameter, minor diameter, rind thickness, rind penetration resistance, and moment of inertia, exhibited significant differences in mean phenotypic values across internodes, highlighting the spatial heterogeneity of geometric and structural determinants of stalk lodging resistance. These findings underscore the importance of internode position in shaping the biomechanical properties that contribute to stalk strength. Variance partitioning revealed that genetic effects were the major contributors to phenotypic variation for most intermediate traits (Figure 1B). However, stalk bending strength was predominantly influenced by environmental variation, suggesting that tissue strength is more susceptible to environmental effects than tissue stiffness, as reflected by the larger genetic component underlying flexural stiffness. Conversely, rind penetration resistance was least affected by environment, suggesting greater phenotype stability and potential for genetic improvement through selection. Along the stalk, the contribution of environment to phenotypic variation increased from the bottom (B1) to the ear (E) internode for all internode traits except rind thickness (Figure 1B). This pattern is consistent with spatial variation in wind loading and the associated mechanosensitive responses along the stalks, which may influence tissue composition, microstructure, and cross-sectional morphology during development, ultimately modulating stalk geometry and structure.

**Figure 1:**
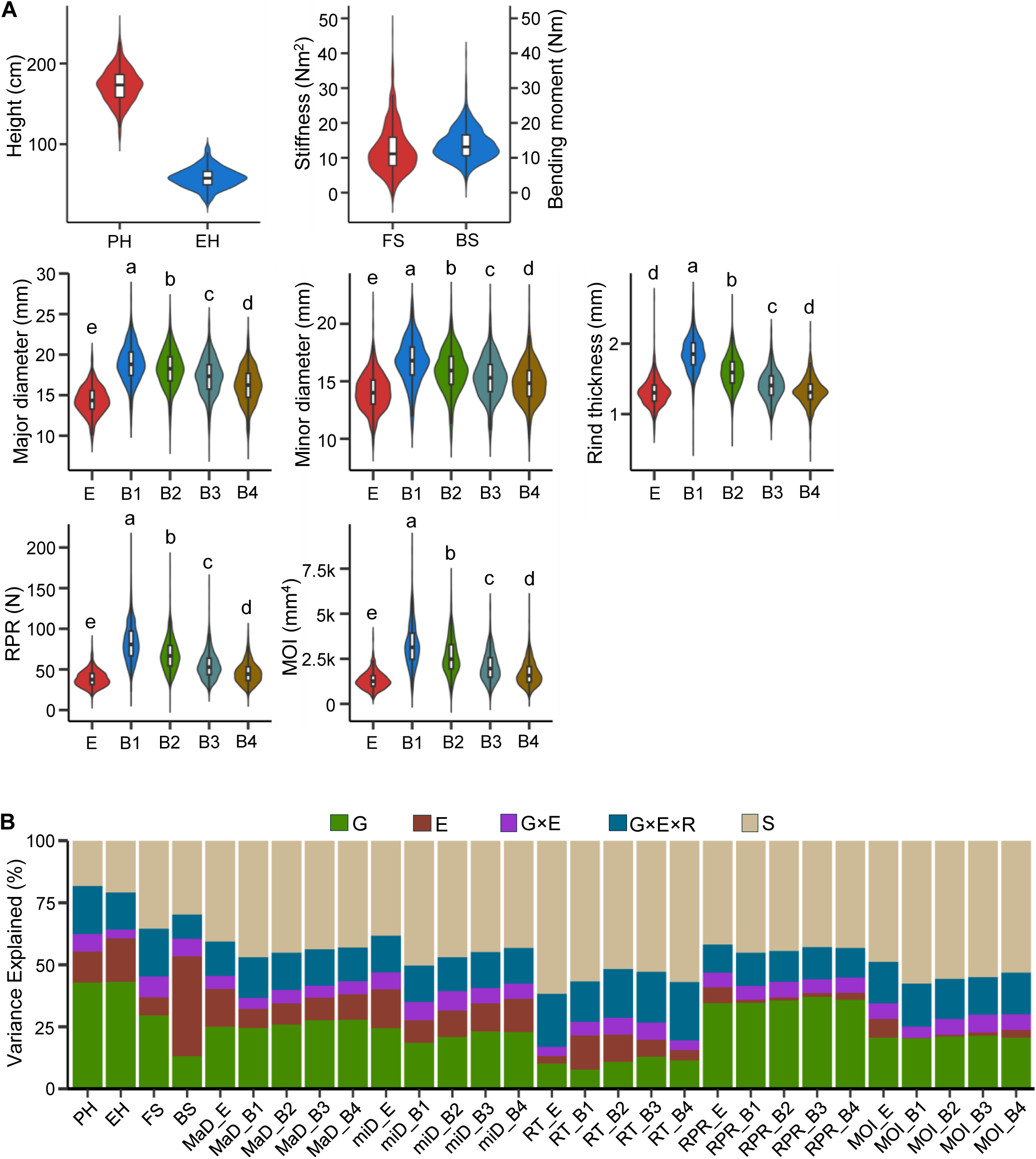
Phenotypic variation and variance partitioning of intermediate traits associated with stalk lodging resistance in maize. (A) Violin plots of adjusted genotype means with embedded boxplots showing the first and third quartiles, and the median represented by a black horizontal line. The x-axis represents the trait or internode names, while the y-axis represents the corresponding phenotypic values. For MOI, the y-axis values are expressed as multiples of k (k = 1000). Different letters (a–e) displayed on the violin plots for the internode traits indicate statistically significant differences among groups, as determined by the Games–Howell test. (B) Stacked bars illustrate the proportion of total variance in each intermediate trait explained by six modeled factors. MaD, Major diameter; miD, Minor diameter; RT, Rind thickness; RPR, Rind penetration resistance; MOI, Moment of inertia; E, Ear internode; B1, Bottom internode; B2–B4, Elongated internodes above B1; G, Genotype; E, Environment; G×E, Genotype by environment interaction; G×E×R, three-way interaction between genotype, environment, and replication; S, Residual.

Variance partitioning of the intermediate traits revealed a substantial genotype-by-environment (G×E) interaction component, indicating that genetic effects were strongly influenced by environment (Figure 1B). Accordingly, we compared the performance of individual inbred lines across environments, revealing phenotypic plasticity and substantial variation in plasticity among inbred lines for the intermediate traits (Figure 2A, Figure S3). The proportion of inbred lines with plastic responses ranged from 89 to 99% for stalk traits and 29.9 to 88.4% for internode traits (Table S2). Only about 3% of the inbred lines exhibited global plasticity across all intermediate traits indicating high sensitivity to environmental variations. Plasticity was more prevalent for stalk traits, indicated by a larger proportion of inbred lines exhibiting plastic responses as compared to internode traits. Remarkably, plasticity for stalk bending strength was nearly universal, with 99% of inbred lines exhibiting differential phenotypic responses along environmental gradients. Apart from G×E, variance partitioning also revealed a substantial G×E×R component underpinning all intermediate traits, suggesting an important role of microenvironment and management practices in driving phenotypic variation.

**Figure 2:**
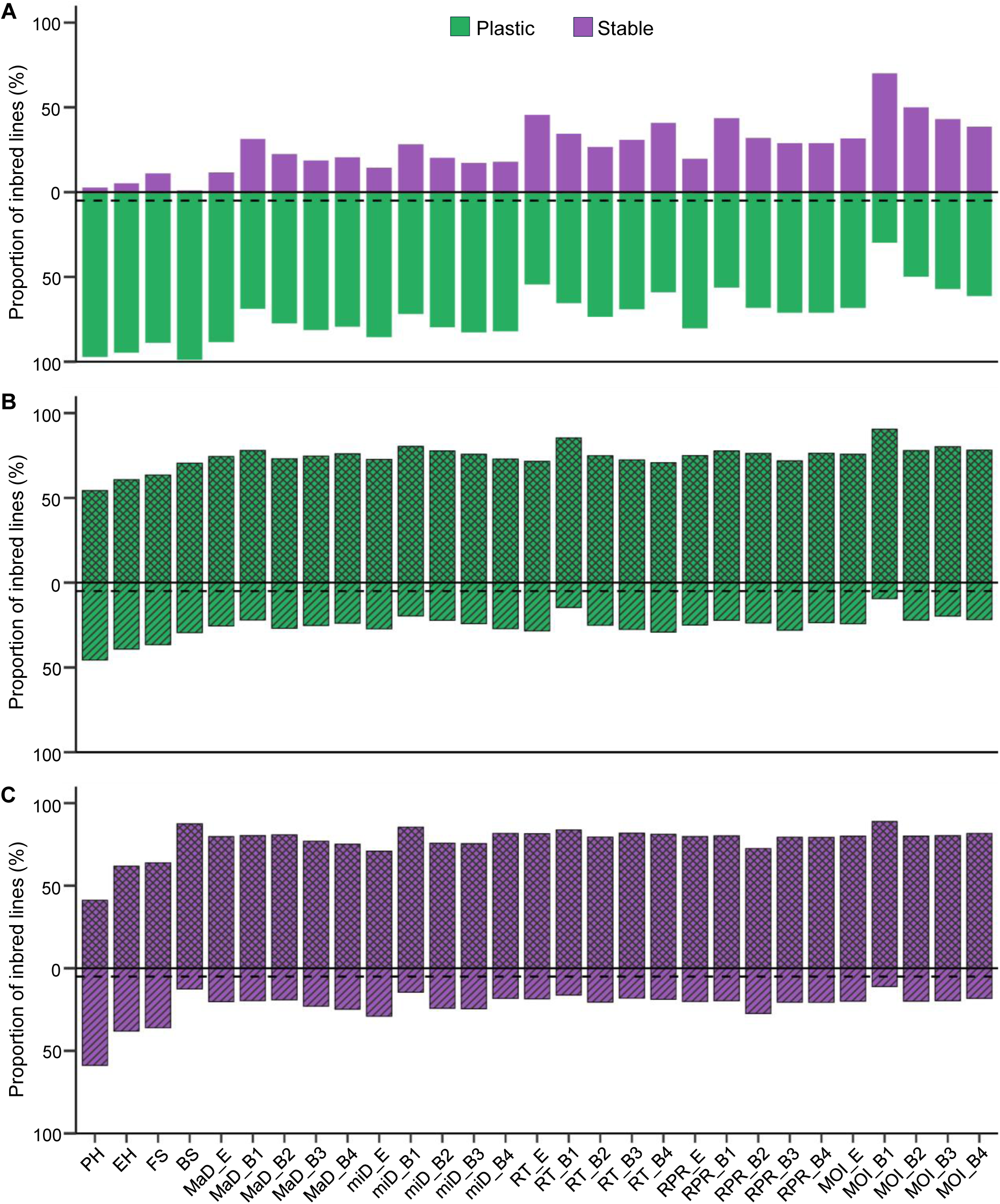
Variation among inbred lines in phenotypic plasticity for intermediate traits associated with stalk lodging resistance. (A) For each trait, the performance of individual inbred lines was compared across environments. Lines exhibiting significant phenotypic differences among environments were classified as plastic and the rest as stable. Plastic and stable classes are shown in emerald green and amethyst purple, respectively, and are separated by a solid horizontal line. The dashed horizontal line indicates α = 0.05. (B-C) Within each environment, replicate means were compared for plastic (B) and stable (C) inbred lines. Lines exhibiting significant replicate differences were sensitive to microenvironment (striped pattern) while the rest were non-sensitive (crosshatched pattern). Abbreviations for intermediate traits and internodes are as defined in Figure 1.

To parse micro-and macroenvironmental drivers of plasticity, we examined the replicate means of intermediate traits within each environment for inbred lines classified as plastic or stable. Within the plastic group, replicate means were not significantly different for more than 50% of inbred lines across all traits (Figure 2B). Despite the substantial G×E×R component identified by variance partitioning, the absence of significant differences among replicates for most inbred lines suggests that replicate-level variation had limited effects on mean phenotypic performance for most genotypes. However, the remaining lines exhibited significant differences among replicates, indicating greater sensitivity to replicate-specific environmental variation. Stalk traits, particularly plant height, showed higher microenvironmental sensitivity, whereas traits measured on individual internodes were less sensitive, suggesting a conserved biological strategy to support stalk structural integrity. Within the stable group, replicate-level patterns were generally consistent with those observed in the plastic group. Although a subset of inbred lines exhibited sensitivity to replicate-specific conditions, these effects were largely offset across environments, yielding overall stability (Figure 2C). Finally, for most traits, the phenotypic distributions of inbred lines exhibiting consistent plasticity or stability across and within environments showed sufficient variation and were broadly representative of the overall inbred panel, indicating that these subsets were suitable for subsequent analyses and unlikely to be strongly affected by sampling bias (Figure S4). Collectively, these findings characterize natural variation and phenotype plasticity for intermediate traits associated with stalk lodging resistance and define plastic and stable genotypes that can be leveraged for subsequent mapping of G×E and gene discovery for lodging-related intermediate traits.

### Multivariate analysis and genetic tractability of intermediate traits

We performed principal component analysis to reduce the dimensionality of the intermediate traits and characterize their relationships. The first five principal components (PC) explained 78.96% of the variance for intermediate traits (Figure 3A, Table S3). All intermediate traits exhibited positive loadings on PC1 ranging from 0.17 to 0.91, which accounted for 47.73% of the total variance. PC1 captured the overall stalk robustness, with higher PC1 scores corresponding to higher stalk flexural stiffness, stalk bending strength, major diameter, minor diameter, moment of inertia, rind thickness, and rind penetration resistance (Figure 3B). Notably, stalk flexural stiffness, stalk bending strength, major diameter, minor diameter, and moment of inertia were tightly clustered indicating high correlation and shared biological mechanisms. PC2 explained 13% of the variance and contrasted between the rind properties and cross-sectional geometry of internodes. Rind thickness and rind penetration resistance showed positive loadings ranging from 0.36 to 0.71, while major diameter and minor diameter had negative loadings from-0.31 to-0.1 (Figure 3B). Higher PC2 scores, therefore, represent thicker and stronger rind with thinner stalks. This pattern suggests a structural tradeoff in stalk architecture wherein resources are differentially partitioned between ground tissue (e.g., larger pith) and mechanical tissue (e.g., thicker or denser rind), thus unfolding diverse but integrated architectural pathways underlying stalk lodging resistance. The remaining PCs each explained a smaller proportion (<10%) of the variance (Figure 3A).

**Figure 3:**
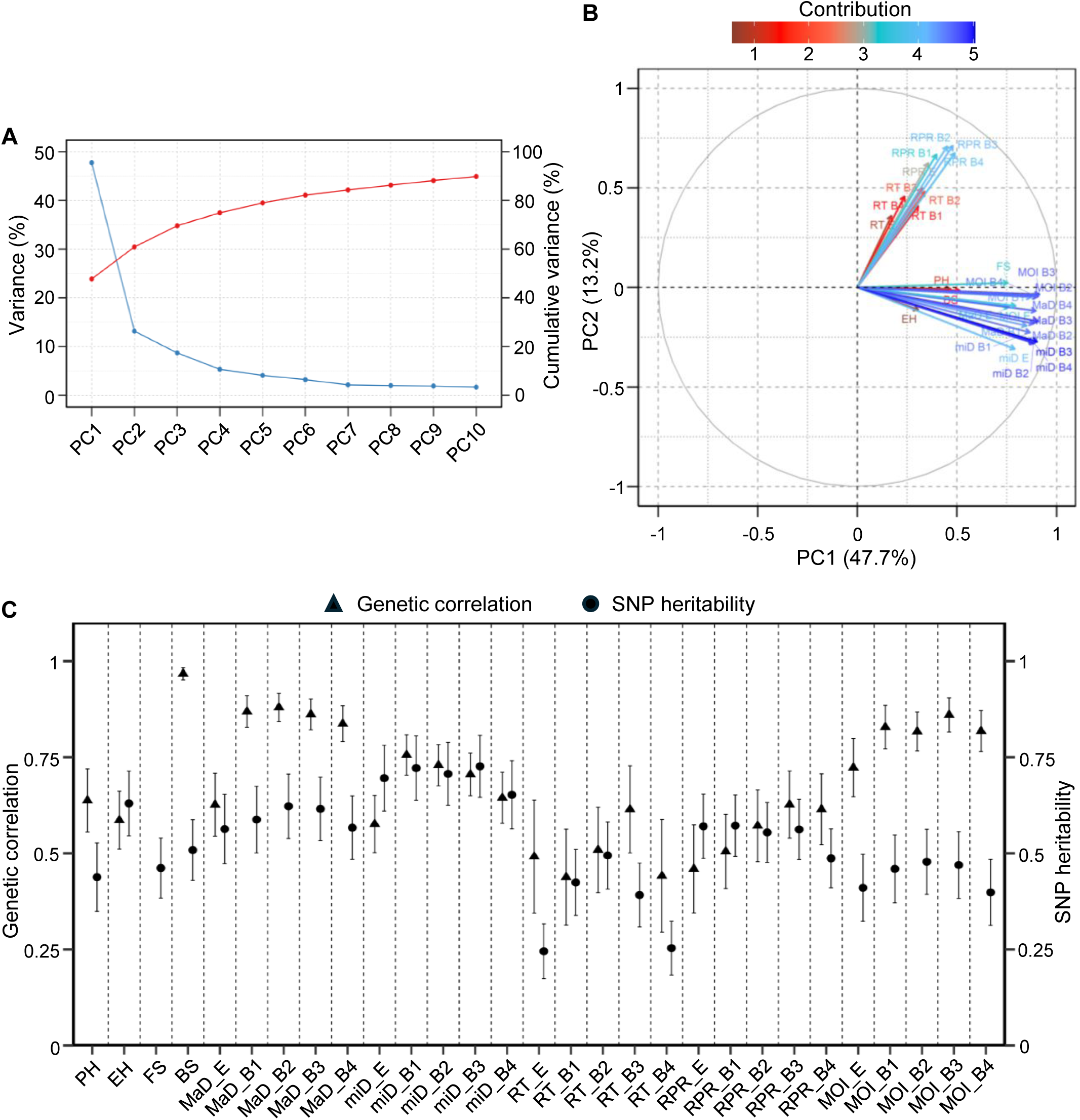
Multivariate analysis of intermediate traits. (A) Proportion of total phenotypic variance explained by the first 10 principal components. The blue line represents variance explained by each principal component (PC), and the red line represents cumulative variance explained. (B) Biplot of loadings for the first two PCs. Colored arrows indicate trait loadings, with arrow length proportional to loading magnitude. PC1 and PC2 are shown on the x-and y-axes, respectively. (C) Genetic correlations and SNP heritability of intermediate traits. Traits are listed on the x-axis and estimates of correlation and heritability are shown on the y-axis. Error bars indicate standard errors. Abbreviations for intermediate traits and internodes are as defined in Figure 1.

To assess the effectiveness of genetic selection for intermediate traits in improving stalk lodging resistance, we estimated SNP heritability (*h_SNP_*^2^), a genetic parameter representing the proportion of variance in a phenotype explained by all common SNPs (Figure 3C). The *h_SNP_*^2^ estimates for stalk and internode traits ranged from 0.44 (SE = 0.09) to 0.63 (SE = 0.08) and 0.25 (SE = 0.07) to 0.73 (SE = 0.08), respectively. The common variants analyzed in the present study explained about 40 to 70% of the total phenotypic variance in most intermediate traits, suggesting strong additive genetic control. Particularly, substantial additive variance (*h_SNP_*^2^>0.55) for ear height, minor diameter, major diameter, and rind penetration resistance indicate effective selection response and enhanced prediction accuracy for these traits. Contrastingly, rind thickness exhibited lower and more variable *h_SNP_*^2^ estimates across internodes compared with other traits, implying a larger role of non-additive effects and limited genetic tractability through selection.

To infer the correlated response to selection in stalk flexural stiffness, a reliable and non-destructive proxy for stalk lodging resistance, we estimated genetic correlations (*r_G_*) with other intermediate traits (Figure 3C). The *r_G_* estimates of stalk flexural stiffness with stalk and internode traits ranged from 0.59 (SE = 0.08) to 0.97 (SE = 0.02) and 0.44 (SE = 0.12) to 0.88 (SE = 0.04), respectively. Consistent with previous observations (Robertson et al., 2016; Tabaracci et al., 2024), stalk bending strength, a destructive trait, showed a very high (*r_G_* > 0.9) correlation with flexural stiffness demonstrating a highly overlapped genetic control. Interestingly, plant height showed high (0.6 < *r_g_* < 0.9) correlation with stalk flexural stiffness indicating shared genetic control and a correlated response to selection. Among the internode traits, major diameter, minor diameter, and moment of inertia exhibited high correlations with stalk flexural stiffness, highlighting substantial pleiotropy between stalk geometry and flexural stiffness. Both rind traits, including rind penetration resistance and rind thickness, showed moderate (0.3 < *r_g_* < 0.6) to high *r_G_* estimates indicating lower correlated response for stalk flexural stiffness as compared to the geometric traits. Phenotypic correlations among intermediate traits mirrored the patterns observed for genetic correlations and supported the findings of our genetic analyses (Figure S5). Altogether, these results identify intermediate traits, particularly internode diameter, moment of inertia, and rind penetration resistance as informative selection indices for improving stalk strength and as entry points for enhancing the genetic resolution of stalk lodging resistance.

### Predictive analytics of stalk lodging resistance

To identify the most influential drivers of stalk lodging resistance, we employed machine learning–based regression to model stalk flexural stiffness and stalk bending strength. XGboost model trained solely on phenotypic data identified major diameter of B1, plant height, and moment of inertia of B2 as the strongest predictors of stalk flexural stiffness (Figure 4A), while the major and minor diameter of B1 and ear height emerged as the most informative predictors of stalk bending strength (Figure 4B). The identification of the major diameter of B1 as the strongest predictor of both flexural stiffness and bending strength reinforces the pivotal role of bottom internode in determining stalk structural integrity and mechanical strength. The model, evaluated using the correlation coefficient (*r*) and root mean squared error (RMSE), exhibited strong and highly significant (P < 0.001) agreement between predicted and observed values. For stalk flexural stiffness, the model achieved *r* value of 0.85 and 0.81 and RMSE value of 5.61 and 6.25 in training and test datasets, respectively. For stalk bending strength, model had r values of 0.70 and 0.60 and RMSE values of 7.80 and 8.58 in training and test sets, respectively. These findings highlight that, besides plant and ear height, stalk geometry, especially internode diameter and moment of inertia, have substantial influence on stalk strength and provide a data-driven basis for trait prioritization in phenotyping and selection.

**Figure 4:**
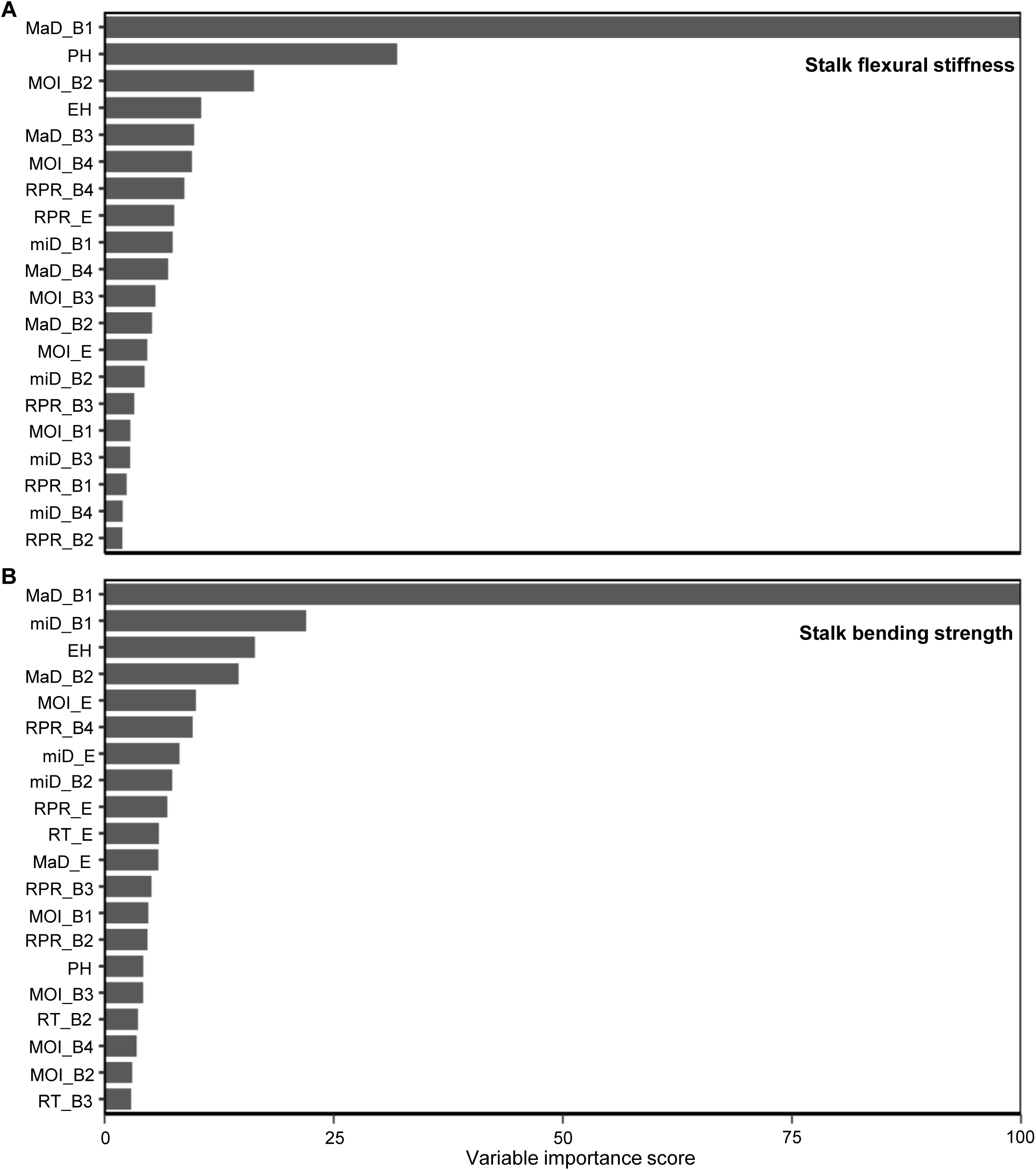
Variable importance of intermediate traits for prediction of stalk flexural stiffness and stalk bending strength. Variable importance scores for intermediate traits are indicated on the x-axis, and the top 20 intermediate traits are listed on the y-axis in decreasing order of their scores. Abbreviations for intermediate traits and internodes are as defined in Figure 1.

### Genetic architecture of intermediate traits

To uncover the loci underpinning intermediate traits, we performed GWA analyses using adjusted means. Bootstrap resampling association analyses with the FarmCPU model yielded 16,371 significant marker-trait associations (Figure 5A, Table S4). For downstream analyses, we filtered these marker-trait associations based on RMIP threshold (see methods) and focused on 705 SNPs, corresponding to 1,001 associations. Moment of inertia of B2 and rind penetration resistance of B4 each contributed the largest proportion of associations (4.30%), whereas rind thickness of B4 accounted for the smallest proportion (2.18%) (Table S5). Major diameter, minor diameter, and moment of inertia across the B1, B2, and B3 internodes collectively accounted for nearly one-third (31.77%) of all marker-trait associations analyzed, reflecting the strong genetic basis of variation in stalk geometry. Shared loci among internodes for an intermediate trait further suggest common genetic control, while differences in gene regulation and environmental effects likely contribute to structural variation among internodes (Figure 5B). Notably, only 19.72% and 4.96% of the 705 variants overlapped genic regions and coding sequences in the maize genome, respectively, underscoring the predominant role of noncoding variation (Table S6). To assess the variant effects of the 705 significant SNPs, we examined their zero-shot scores generated by PlantCaduceus, a plant DNA language model (Zhai et al., 2025). The zero-shot scores ranged from −5.97 to 6.70, with more than 54% of the SNPs exhibiting negative scores, highlighting an enrichment of variants located in genomic regions under greater sequence constraint (Figure S6).

**Figure 5:**
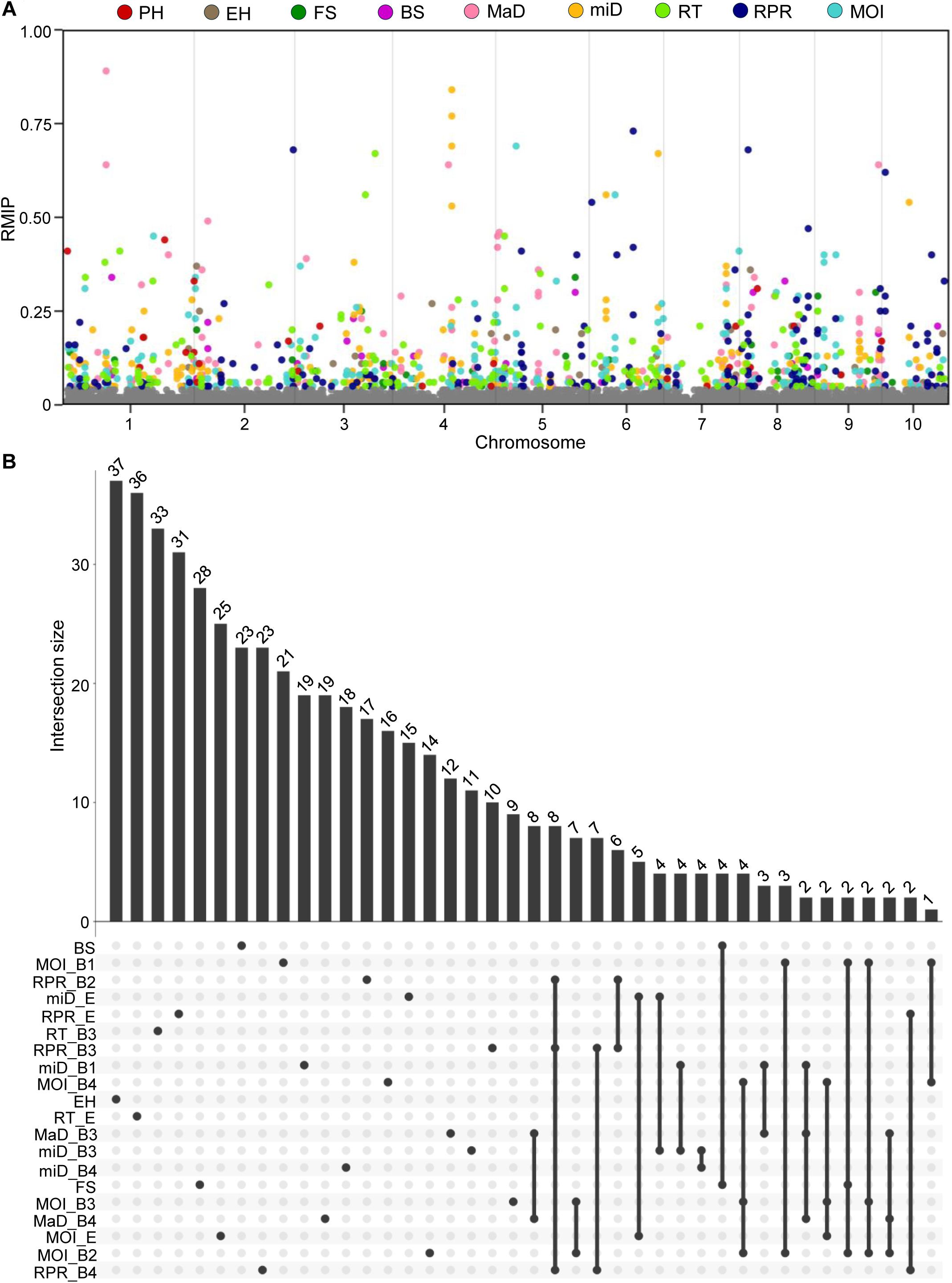
Marker-trait associations for intermediate traits (A) Manhattan plot showing significant SNPs associated with intermediate traits. For each trait, SNPs with RMIP scores ≥ 0.05 are color coded as shown in the legend, whereas the remaining SNPs are displayed in grey. (B) UpSet plot illustrating the distribution of SNPs (RMIP ≥ 0.05) underlying significant marker-trait associations across intermediate traits, with the 20 most frequent SNP sets displayed. Abbreviations for intermediate traits, internodes, and environments are as defined in Figure 1.

Remarkably, about one-fifth (21.98%) of the 705 variants were associated with at least two traits, highlighting the substantial role of pleiotropy in regulating intermediate traits (Table S6). The three most highly pleiotropic SNPs (chr7_151386854, chr4_143279823, and chr6_40863789) were exclusively associated with stalk geometry traits, each exhibiting 9 to 12 unique marker-trait associations (Table S6). However, all three variants were confined to noncoding regions of the genome and likely represent distal cis-regulatory variation. The most pleiotropic SNP, chr7_151386854, was situated about 8.9 kb upstream of Zm00001eb320310 (*dek47*), while chr4_143279823 and chr6_40863789 were located approximately 3.8 kb and 13 kb upstream of Zm00001eb184630 and Zm00001eb266050, respectively (Figure 6A). Analysis of allelic composition of these loci revealed significant effects on the phenotypic outcome of most intermediate traits. While the rare allele (A) at chr7_151386854 was associated with superior trait performance, the major allele (G) was favorable at chr4_143279823, highlighting contrasting allelic effects between the two loci (Figure 6B, Figure S7-S8).

**Figure 6:**
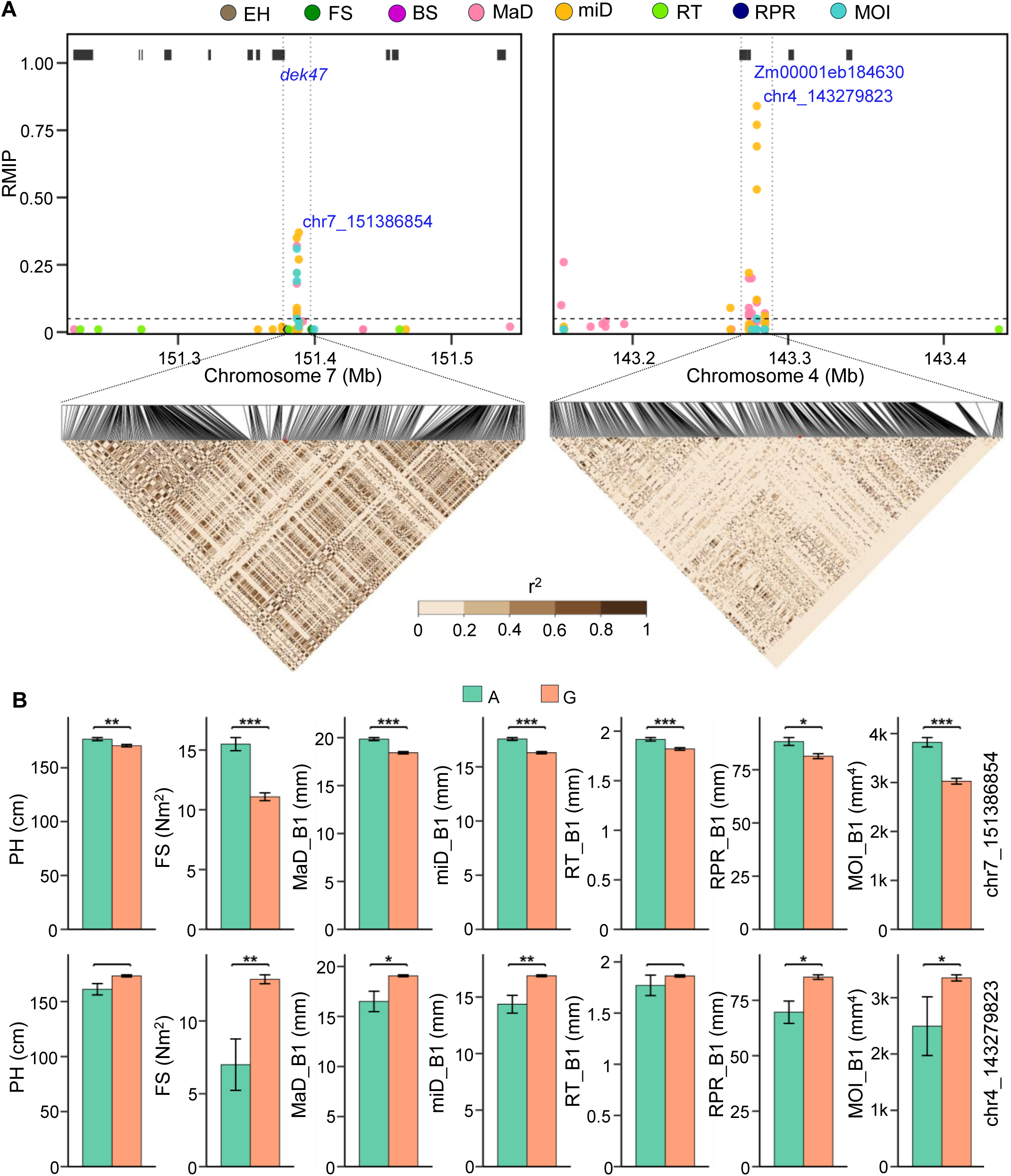
Effects of allele composition at the pleiotropic SNPs chr7_151386854 and chr4_143279823 on phenotype variation in intermediate traits. (A) Local manhattan plots illustrate the distribution of significant SNPs within a 200 kb window surrounding chr7_151386854 and chr4_143279823. Heatmaps show linkage disequilibrium (r^2^) within a 10 kb window around each SNP. Solid black rectangles at the top of each manhattan plot represent annotated genes (B) For each trait, bar plots show mean phenotypic values, with the corresponding values indicated on the y-axis. Error bars represent standard errors, and bar colors indicate SNP genotypes as defined in the legend. Within each bar plot, asterisks denote significant allelic effects, with one, two, and three asterisks indicating P < 0.05, P < 0.01, and P < 0.001, respectively. Abbreviations are as defined in Figure 1.

To elucidate the genetic determinants of intermediate traits, we surveyed ±10 kb genomic intervals surrounding the 705 SNPs and identified 533 unique candidate genes (Table S7). These candidate genes represented 187 superfamilies, with Protein kinase-like (SSF56112), P-loop NTPase (SSF52540), and RING/U-box (SSF57850) being among the most frequently represented and collectively accounting for 12.38% of the candidate genes (Table 1). Among these candidates, several promising genes emerged, including Zm00001eb426640 (*nactf65*) and Zm00001eb367300 (*myb79*), both of which have been associated with rind penetration resistance and reported to regulate lignin accumulation and mechanical strength in leaf tissues (Shi et al., 2026). Interestingly, Zm00001eb143080 (*mads69*) and Zm00001eb353250 (*pebp8*), both of which are involved in the regulation of flowering, have been associated with stalk diameter and moment of inertia, suggesting a potential link between flowering time and stalk geometry. In addition to novel candidates, we recovered genes previously linked to traits associated with stalk lodging resistance in maize. For example, Zm00001eb400780 (*xyl3*) was linked to stalk diameter (Xu et al., 2023) and Zm00001eb414390 (*xth1*), associated with rind penetration resistance, showed differential expression between inbred lines B73 and QY1 that differ in stalk mechanical strength (Yang et al., 2024). Collectively, the novel candidate genes open new opportunities to enhance the genetic resolution of stalk lodging resistance, while rediscovered loci validate the biological relevance of the marker–trait associations identified in the present study.

**Table 1:** List of 25 largest gene superfamilies associated with intermediate traits.

| **Accession** | **No. of candidate genes** | **Superfamily name** |
| --- | --- | --- |
| SSF56112 | 26 | Protein kinase-like (PK-like) |
| SSF52540 | 23 | P-loop containing nucleoside triphosphate hydrolases |
| SSF57850 | 17 | RING/U-box |
| SSF46689 | 13 | Homeodomain-like |
| SSF51735 | 13 | NAD(P)-binding Rossmann-fold domains |
| SSF48371 | 9 | ARM repeat |
| SSF54171 | 9 | DNA-binding domain |
| SSF52047 | 7 | RNI-like |
| SSF52058 | 7 | L domain-like |
| SSF53474 | 7 | alpha/beta-Hydrolases |
| SSF46565 | 6 | Chaperone J-domain |
| SSF51445 | 6 | (Trans)glycosidases |
| SSF53335 | 6 | S-adenosyl-L-methionine-dependent methyltransferases |
| SSF49785 | 5 | Galactose-binding domain-like |
| SSF53756 | 5 | UDP-Glycosyltransferase/glycogen phosphorylase |
| SSF54928 | 5 | RNA-binding domain, RBD |
| SSF47459 | 4 | HLH, helix-loop-helix DNA-binding domain |
| SSF48264 | 4 | Cytochrome P450 |
| SSF49493 | 4 | HSP40/DnaJ peptide-binding domain |
| SSF49899 | 4 | Concanavalin A-like lectins/glucanases |
| SSF50978 | 4 | WD40 repeat-like |
| SSF53383 | 4 | PLP-dependent transferases |
| SSF56235 | 4 | N-terminal nucleophile aminohydrolases (Ntn hydrolases) |
| SSF56784 | 4 | HAD-like |
| SSF57959 | 4 | Leucine zipper domain |

## Discussion

### Phenotype plasticity varies along the stalk and is primarily driven by macroenvironment

The relative contributions of micro-and macroenvironments to phenotypic variation and plasticity for complex traits, including those related to stalk lodging resistance, remain poorly understood. Our study indicates that plastic responses are largely determined by macroenvironmental differences while microenvironmental sensitivity exists in a smaller subset of the sampled germplasm. The existence of such variation provides opportunities to map underlying loci and, where desirable, to select genotypes that are resistant to small-scale environmental variations. We also observed a clear spatial gradient for phenotype plasticity along the stalk, as trait sensitivity to environmental effects increased from basal to upper internodes. Wind loading typically varies along the crop sub-canopy, generating an internal bending moment that increases basipetally along the stalk. However, the larger bending moments experienced by basal internodes are offset by their increased diameters, resulting in a relatively uniform distribution of mechanical stress along the stalk. In contrast, flexural deflection under wind loading increases progressively from basal to apical internodes, resulting in greater mechanical displacement in upper internodes. This deflection gradient may elicit differential mechanosensory responses that contribute to the distinct patterns of phenotype plasticity observed among internode properties. While internode morphology is influenced by localized microenvironmental variables at the corresponding sub-canopy layer, stalk traits represent outcomes of carbon partitioning, biomass accumulation, and growth dynamics along entire sub-canopy. As a result, stalk traits capture cumulative developmental responses to environmental variations, including changes in light availability, water status, wind exposure, etc., and therefore, exhibit stronger sensitivity to environmental variation than internode traits.

Emergence of macroenvironment as the primary driver of phenotype plasticity has key implications in breeding for stalk lodging resistance. For example, in multi-location trials, breeders can exploit plasticity to select genotypes with specific adaptability for particular environments, delivering predictable and stable crop performance in such conditions (Ceccarelli and Grando, 2020). However, broad spatial gradients in environmental conditions often trigger crossover G×E, where genotype rankings shift across environments, compromising selection efficiency for genotypes exhibiting homeostasis across diverse environments (Cooper and Podlich, 1999). Therefore, selection models that explicitly account for heterogeneity in phenotype plasticity and G×E in multilocation breeding programs can separate stable genetic effects from environment-specific responses, enabling accurate germplasm evaluation in crop improvement programs (Malosetti et al., 2013; Rebollo et al., 2023).

### Stalk geometry and rind architecture represent complementary pathways to stalk strength

Comprehensive phenotyping of various intermediate traits revealed that stalk cross-sectional geometry and rind architecture represent two fundamental drivers of stalk lodging resistance (Figure 3B). While cross-sectional geometry, especially internode diameter and moment of inertia, sets the stalk biomechanical leverage against bending, rind strength, reflected in cell wall thickness and penetration resistance, determines tissue toughness. Mechanistically, stalk strength and stiffness increase when both geometric and material properties are enhanced together. For example, flexural stiffness is approximately E x I, where inherent rind properties modulate the material property (E) and cross-sectional geometry determines the second moment of inertia (I) (Robertson et al., 2017; Oduntan et al., 2024). Interestingly, multivariate analysis revealed a trade-off rather than mutual exclusivity between geometry and rind properties, suggesting three mechanistic pathways to achieve stalk strength: 1) thicker or denser rinds at smaller stalk diameters, 2) larger diameters with lesser rind emphasis, and 3) balanced gains in both traits. This interpretation aligns with fundamental cantilever beam mechanics and is consistent with findings from prior field studies linking stalk geometry to lodging risk, as well as the strong correlation between stalk flexural stiffness and bending strength observed across diverse maize genotypes (Robertson et al., 2016; Stubbs et al., 2022). The relevance of three mechanistic pathways for stalk strength is demonstrated by genetic mapping studies that consistently identify signals for both geometric (Bian et al., 2024; Boatwright et al., 2024) and rind traits (Wu et al., 2022) across diverse grass species, including maize.

From a breeding perspective, improving stalk lodging resistance requires a selection index that recognizes the complementary nature of geometric and rind properties, optimally balancing both parameters to enhance stalk strength. Balancing this geometry-rind trade-off is critical, as high rind penetration resistance is often accompanied by elevated lignin content that compromises stover digestibility (Martin et al., 2004), while aggressively increasing stalk robustness can divert resources from grain filling and reduce yields (Zhang et al., 2023). An equally important consideration when deploying such an index is maintaining plant height within desirable limits, as taller plants substantially increase the bending moment and lodging risk (Stubbs et al., 2023). Ultimately, the index weights should be optimized to the breeding goals, end use, expected lodging pressure, planting density, and target environment to enhance stalk lodging resistance while maximizing grain yields.

### Pleiotropy and regulatory variation underpin the genetic basis of intermediate traits

Tight associations of internode diameter, moment of inertia, and rind penetration resistance of basal internodes with stalk flexural stiffness are consistent with shared genetic control, while their genetic tractability indicates that selection for stalk geometry and rind strength can deliver correlated gains in enhanced lodging resistance (Figure 3C, Figure S5). While the role of cross-sectional morphology in determining stalk mechanical performance was previously established, stalk geometry remains an underutilized breeding target for improving stalk lodging resistance in maize (Stubbs et al., 2022). In addition to the correlation analyses, findings from predictive modeling demonstrate significant potential of targeted geometric modifications to enhance stalk strength in maize. These interpretations align with previous reports demonstrating that robust cross-sectional morphology improves stalk mechanical performance (Forell et al., 2015; Zhao et al., 2025). Together, these insights highlight stalk geometry as a key breeding target for improving stalk lodging resistance in maize.

Our study revealed that noncoding variation accounted for a significant proportion of the heritable variance in intermediate traits, as only <5% of the analyzed variants overlapped the coding sequences in the maize reference genome (Table S6). While advances in sequencing and computational tools enabled large-scale assembly of complex genomes, the functional annotation of regulatory variation remains largely unexplored in plants yet is rapidly gaining traction (Song et al., 2021; Marand et al., 2023). Particularly in maize, regulatory variation has been established as a key driver of natural variation for various agronomic traits (Li et al., 2012; Wallace et al., 2014). However, the functional variants underlying phenotypic diversity in stalk lodging resistance remain poorly characterized. Characterization of noncoding variation facilitates identification of cis-regulatory elements for trait improvement by optimizing expression of target protein-coding genes. For example, altering gene expression through promoter modifications in *stiff1* and *CLE* genes significantly enhanced stalk strength and grain-yield-related traits, respectively, in maize (Zhang et al., 2019; Liu et al., 2021).

### Candidate genes reflect regulatory control of intermediate traits

While individual association signals from GWA analyses rarely pinpoint a single causal gene underlying complex plant traits, G×E further affect the consistency of these association signals across diverse environments. Therefore, we prioritized association mapping of intermediate traits using genetic effects estimated across environments to identify global association signals with broad relevance to crop improvement. Analysis of association signals revealed a sizeable proportion of candidate genes encoding transcription factors, consistent with a role for gene regulation in determining variation in intermediate traits, as reported previously (Wallace et al., 2014). The recovery of genes coding for transcription factors previously implicated in traits associated with stalk lodging resistance further supports the validity of our mapping approach. For example, Zm00001eb138920 (*myb8*), which emerged as a candidate gene for stalk bending strength, has been reported to act as an activator of lignin biosynthesis in maize stalks (Zhan et al., 2024). Collectively, these findings provide hypotheses for follow-up studies and reinforce the need for integrating expression and chromatin data, fine-mapping, and functional validation to establish the role of regulatory variation and candidate genes.

In essence, our study demonstrates that stalk lodging resistance is not a single trait, but a coordinated phenotype emerging from the contributions of many intermediate traits. Stalk geometry, rind architecture, and their associated regulatory variation play central roles in determining stalk lodging resistance. While cross-sectional geometry determines the stalk leverage against bending (I), rind architecture governs both material stiffness (E) and tissue resistance to fracture, thereby shaping the flexural stiffness and bending strength of the stalk. Enrichment of noncoding marker-trait associations and substantial pleiotropy observed in the study indicate that the genetic control of stalk lodging resistance is not concentrated in individual genes but rather distributed across regulatory variation that jointly tunes geometric and material properties. Accordingly, our findings shift the genetics of stalk lodging resistance, and plant structural traits more broadly, from identifying individual causal genes toward understanding of the regulatory architecture that shapes and coordinates underlying components such as E and I. The next step is to functionally characterize candidate loci and to establish gene networks linking geometric and material components to flexural stiffness, thereby providing independent validation to the beam theory-based decomposition of flexural stiffness and advancing stalk strength assessment beyond correlated proxy traits. Scaling this approach to large germplasm panels and, ultimately, across crop species could transform stalk lodging resistance from a selection-dependent phenotype into a mechanistically engineered trait.

## Acknowledgements

We sincerely appreciate all the talented undergraduate students and research staff of the Kentucky-Idaho-Clemson Plant Biomechanics Consortium for assistance in field maintenance and phenotypic data collection. We thank the United States Department of Agriculture - National Plant Germplasm System (https://npgsweb.ars-grin.gov/gringlobal/search) for providing seed material of the maize inbred panel.

## CRediT authorship contribution statement

**Bharath Kunduru –** Methodology, Investigation, Data curation, Formal Analysis, Software, Visualization, Validation, Writing – original draft, Writing – review & editing; **Norbert T. Bokros –** Methodology, Investigation, Data curation, Writing – review & editing; **Kaitlin Tabaracci –** Methodology, Investigation, Data curation; **Rohit Kumar –** Methodology, Investigation, Writing – review & editing; **Manwinder S. Brar –** Methodology, Investigation, Writing – review & editing; **Yusuf Oduntan –** Methodology, Investigation; **Christopher J. Stubbs –** Methodology, Investigation; **Caique Machado e Silva –** Methodology, Writing – review & editing; **William C. Bridges –** Methodology; **Ravi V. Mural –** Methodology, Writing – review & editing; **Gota Morota –** Methodology, Writing – review & editing; **Christopher S. McMahan –** Conceptualization, Methodology, Investigation, Formal Analysis, Validation, Supervision, Funding acquisition, Project administration, Writing – review & editing; **Seth DeBolt –** Conceptualization, Methodology, Investigation, Supervision, Resources, Funding acquisition, Project administration, Writing – review & editing; **Daniel J. Robertson –** Conceptualization, Methodology, Investigation, Data curation, Supervision, Resources, Funding acquisition, Project administration, Writing – review & editing; **Rajandeep S. Sekhon –** Conceptualization, Methodology, Investigation, Data curation, Formal analysis, Supervision, Resources, Funding acquisition, Project administration, Writing – original draft, Writing – review & editing

## Data availability

The metadata relevant to the present study are available on Zenodo (Kunduru et al., 2025).

## Disclaimer

Any opinions, findings, and conclusions or recommendations expressed in this material are those of the author(s) and do not necessarily reflect the views of the National Science Foundation.

