## Supplemental Figure for "Phenotypic plasticity, stalk geometry, and noncoding variation underpin stalk lodging resistance in maize"

Figure S1

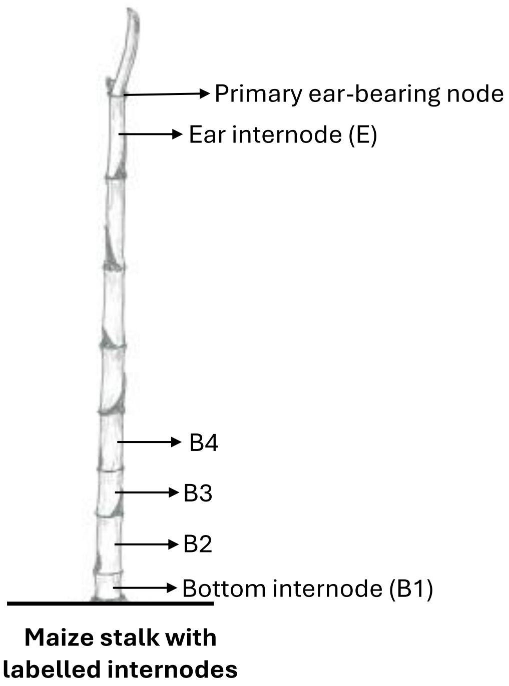


**Figure S1:** Standardized methodology for labelling internodes for phenotyping maize stalks.

Figure S2

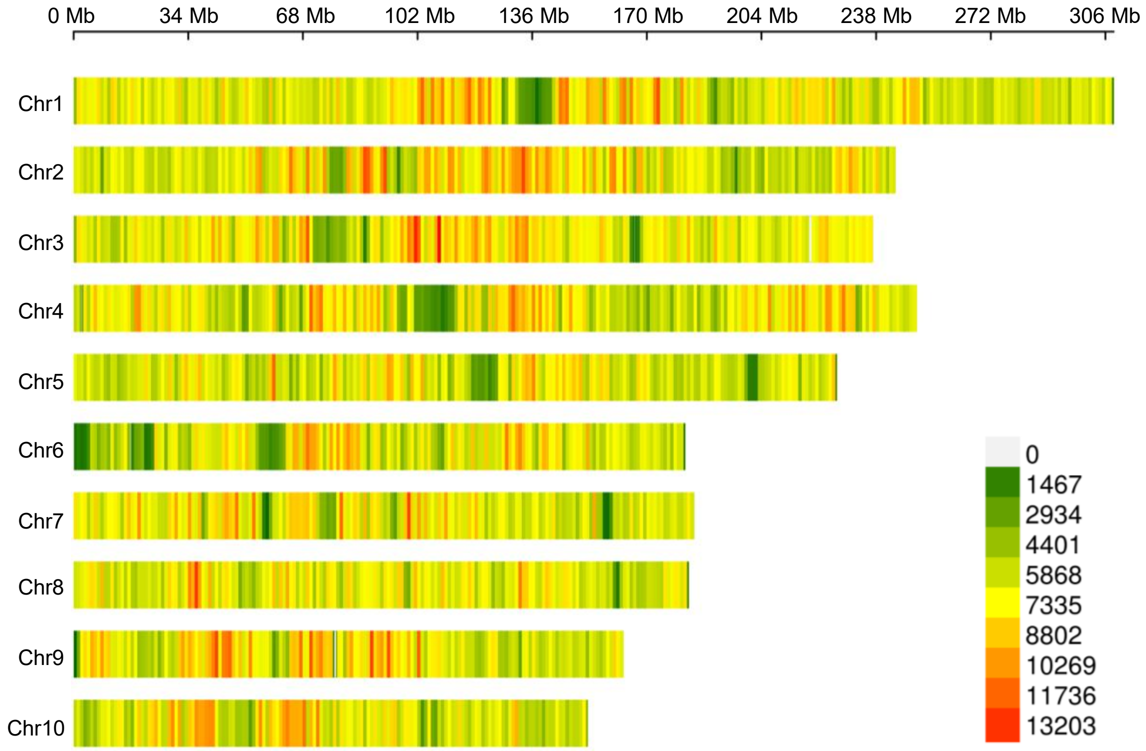


**Figure S2:** Marker density plot showing the distribution of 14,359,923 SNPs across the ten maize chromosomes. Color codes in the legend indicate the number of SNPs.

Figure S3

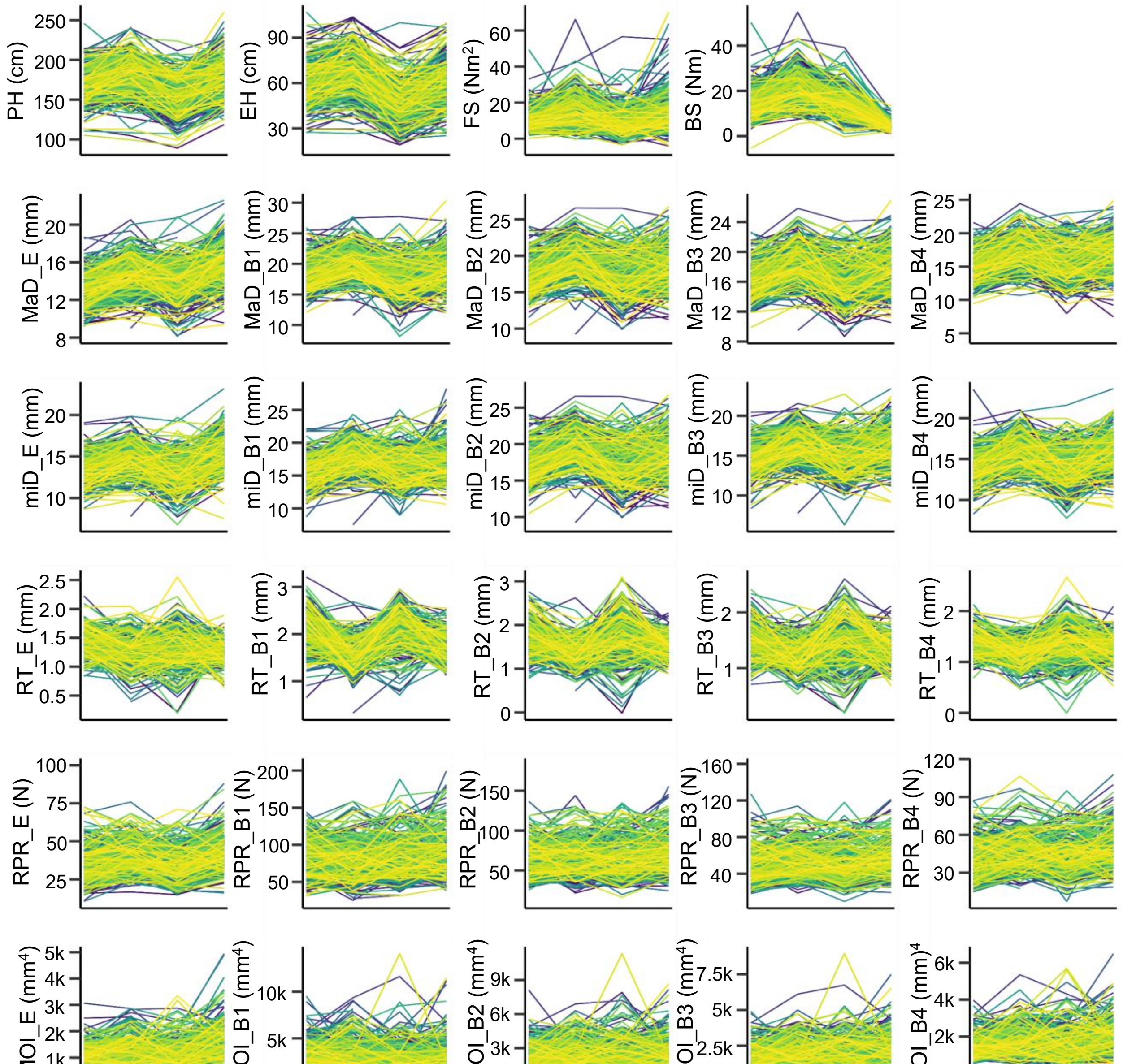


**Figure S3:** Reaction norms of intermediate traits of stalk lodging resistance across different environments. Environments and observed phenotypic values are shown on x- and y-axes, respectively. For each trait, colored lines represent the individual inbred lines studied. MaD, Major diameter; miD, Minor diameter; RT, Rind thickness; RPR, Rind penetration resistance; MOI, Moment of inertia; E, Ear internode; B1, Bottom internode; B2–B4, Elongated internodes above B1; C20, Clemson University 2020; C21, Clemson University 2021; K20, University of Kentucky 2020; K21, University of Kentucky 2021.

Figure S4

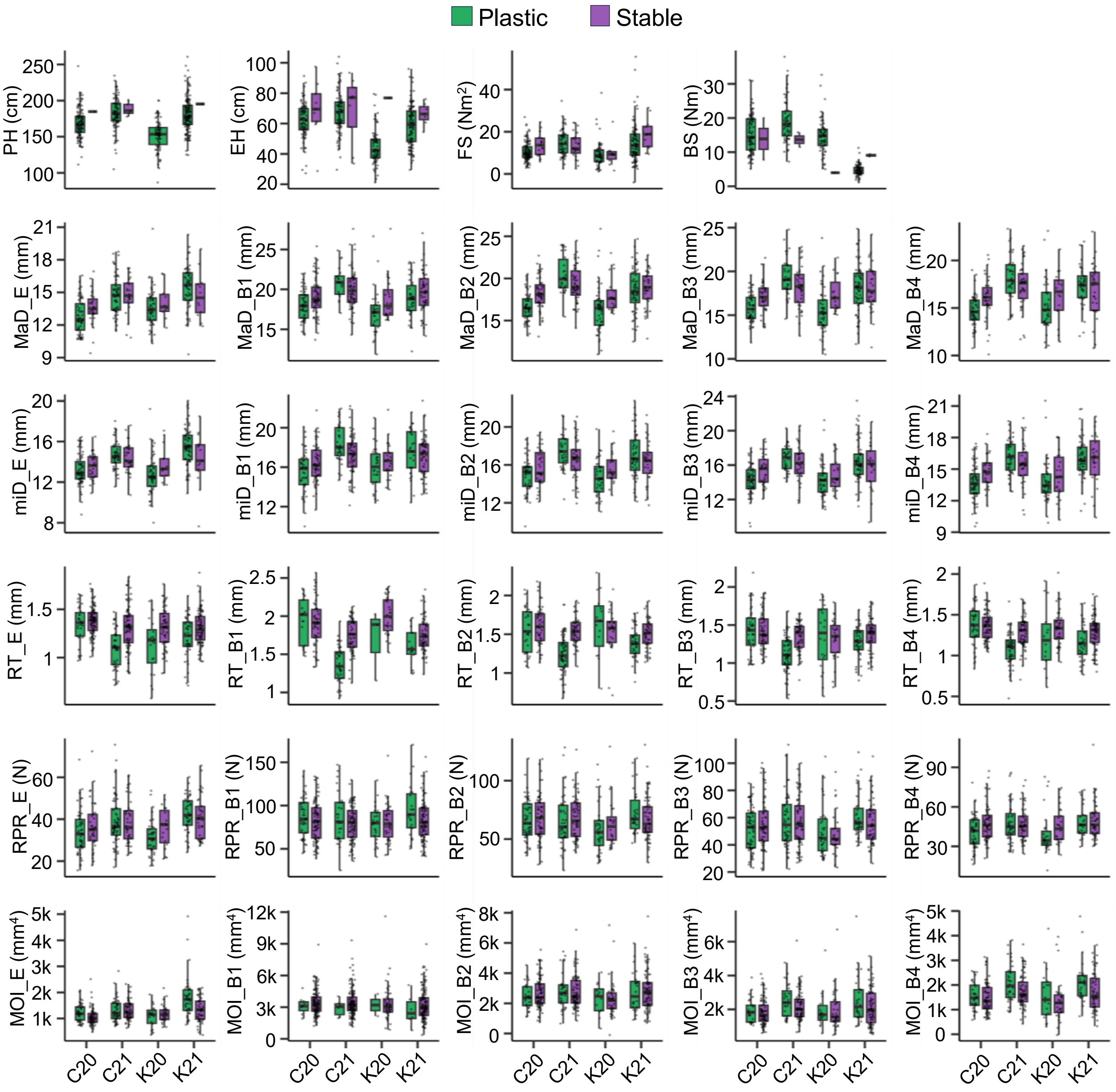


**Figure S4**: Phenotypic distributions of intermediate traits in globally plastic and stable inbred lines. Plastic lines exhibited significant phenotypic variation across and within environments whereas, stable lines showed consistent phenotypic values across and within environments. Abbreviations of intermediate traits, internodes, and environments are as defined in Figure S3.

Figure S5

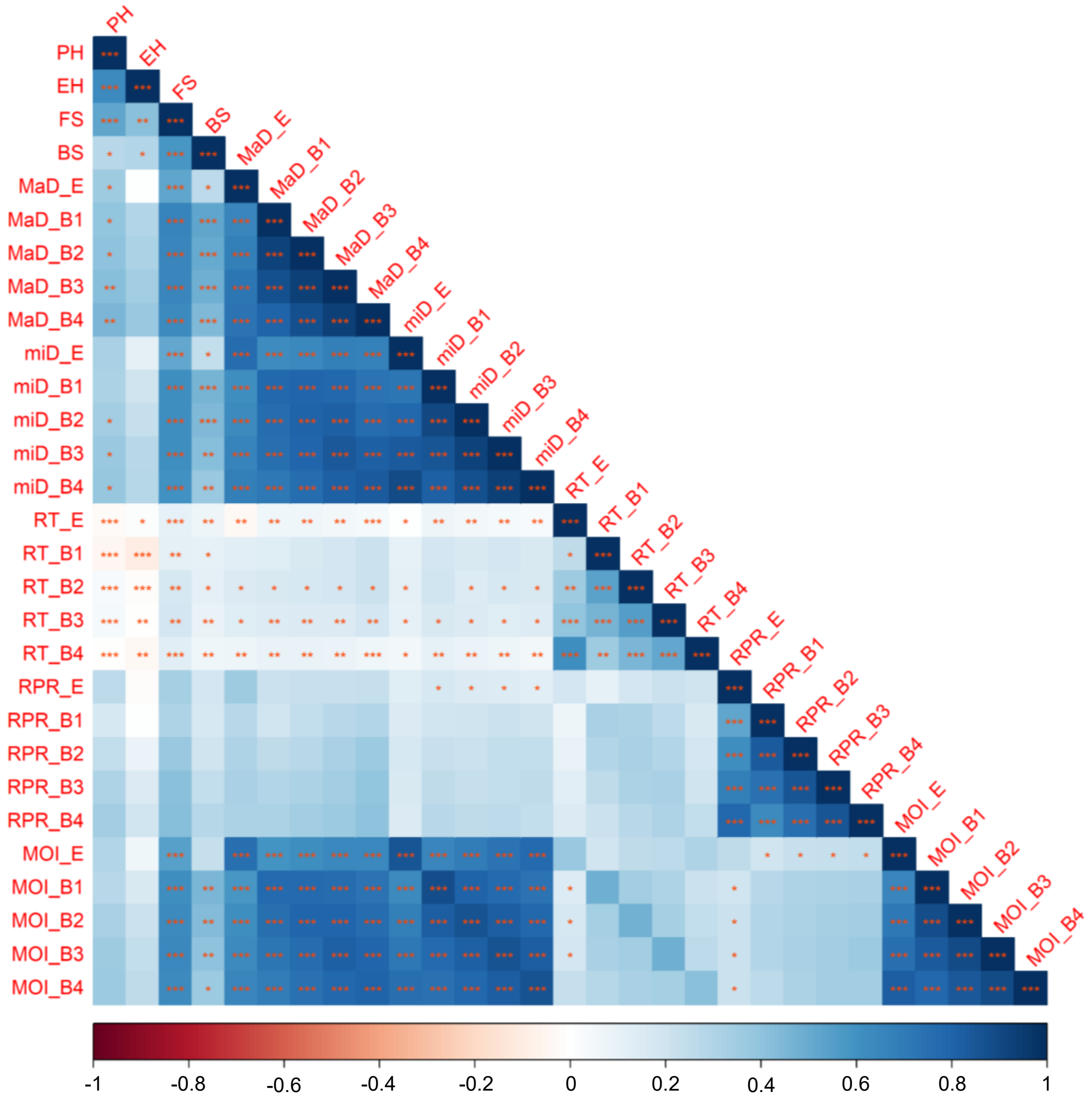


**Figure S5:** Phenotype **c**orrelations among intermediate traits of stalk lodging resistance. The heatmap represents pair-wise Pearson correlation coefficients of traits. Abbreviations are same as defined in Figure S3.

Figure S6

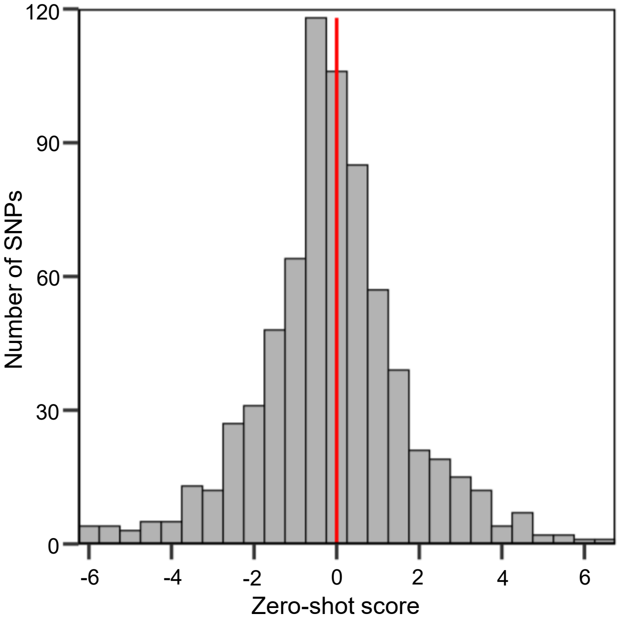


**Figure S6:** Distribution of zero-shot scores associated with significant SNPs (RMIP ≥ 0.05). In the histogram, bars represent the frequency of SNPs within 0.5-unit intervals of zero-shot scores. The red vertical line represents a zero-shot score of 0.

Figure S7

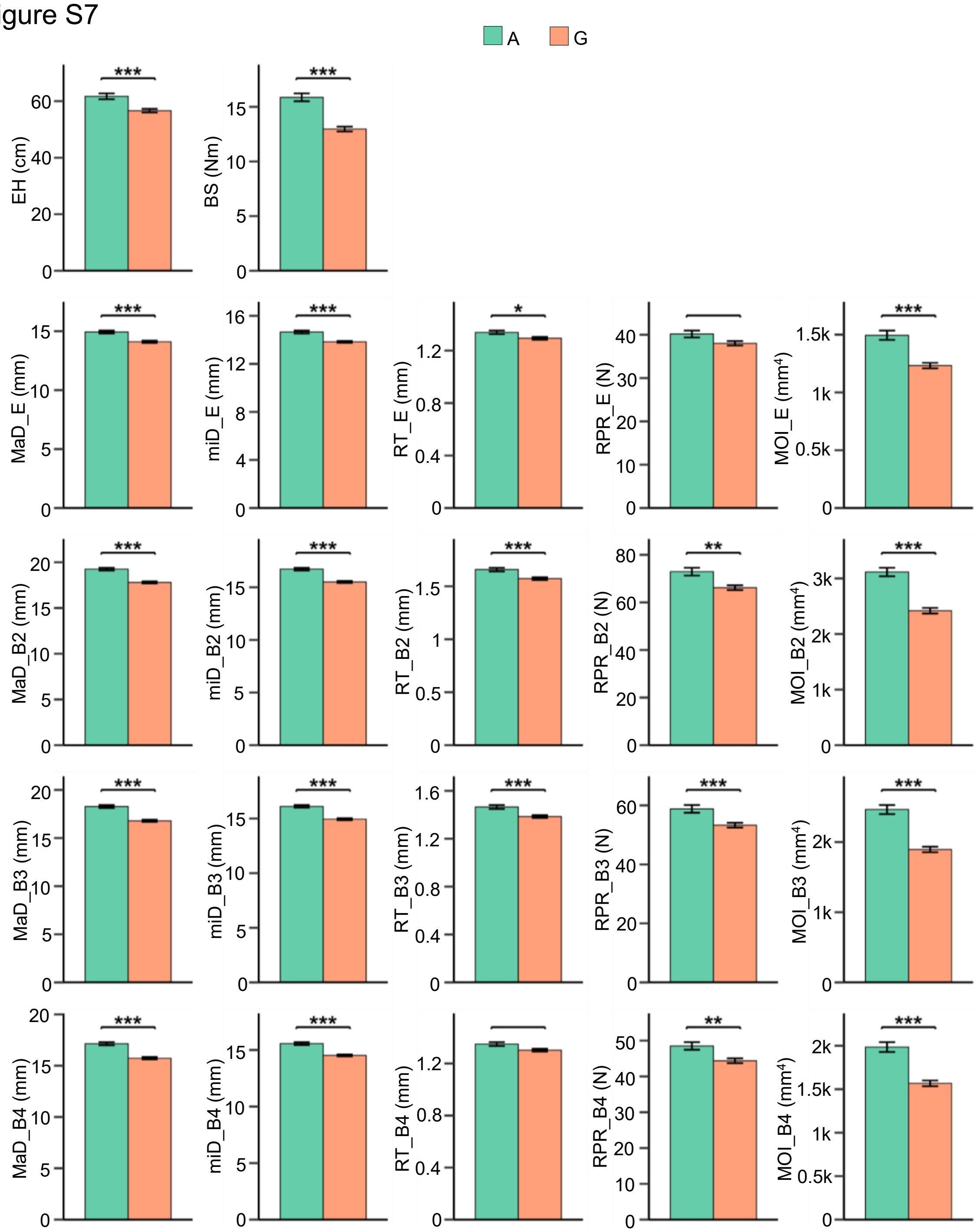


**Figure S7:** Effect of allele composition at SNP chr7_151386854 on phenotype variation in intermediate traits. For each trait, bar plots show mean phenotypic values, with the corresponding values indicated on the y-axis. Error bars represent standard errors, and bar colors indicate SNP genotypes as defined in the legend. Within each bar plot, asterisks denote significant allelic effects, with one, two, and three asterisks indicating $P < 0.05$, $P < 0.01$, and $P < 0.001$, respectively. Abbreviations are same as defined in Figure S3.

Figure S8

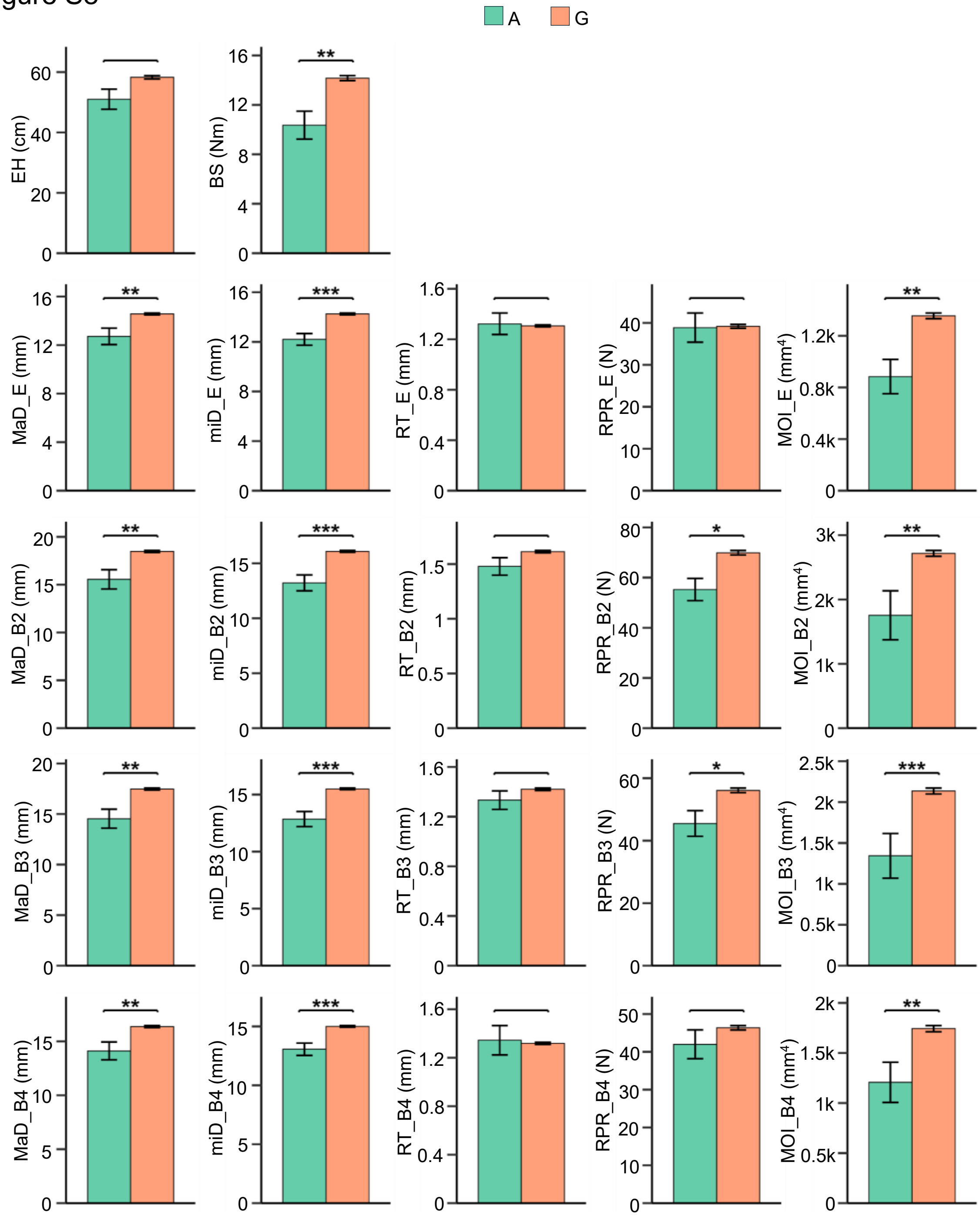


**Figure S8:** Effect of allele composition at SNP chr4_143279823 on phenotype variation in intermediate traits. For each trait, bar plots show mean phenotypic values, with the corresponding values indicated on the y-axis. Error bars represent standard errors, and bar colors indicate SNP genotypes as defined in the legend. Within each bar plot, asterisks denote significant allelic effects, with one, two, and three asterisks indicating $P < 0.05$, $P < 0.01$, and $P < 0.001$, respectively. Abbreviations are same as defined in Figure S3.
